# Evolutionary history of the brown rat: out of southern East Asia and selection

**DOI:** 10.1101/096800

**Authors:** Lin Zeng, Chen Ming, Yan Li, Ling-Yan Su, Yan-Hua Su, Newton O. Otecko, Ambroise Dalecky, Stephen Donnellan, Ken Aplin, Xiao-Hui Liu, Ying Song, Zhi-Bin Zhang, Ali Esmailizadeh, Saeed S. Sohrabi, Hojjat Asadollahpour Nanaei, He-Qun Liu, Ming-Shan Wang, Solimane Ag Atteynine, Gérard Rocamora, Fabrice Brescia, Serge Morand, David M. Irwin, Ming-sheng Peng, Yong-Gang Yao, Hai-Peng Li, Dong-Dong Wu, Ya-Ping Zhang

## Abstract

The brown rat (*Rattus norvegicus*) is found wherever humans live and transmits many diseases, and its breeding produced the laboratory rat used widely in medical research. Here, we sequenced whole genomes from 118 rats to explore the origin and dispersal routes of the brown rat and the domestication of the laboratory rat. We showed that brown rats migrated about 3600 years ago from southern East Asia, rather than Northern Asia as formerly suggested, to the Middle East and then to Europe and Africa. Many genes involved in the immune system experienced positive selection in the wild brown rat, while genes involved in the nervous system and energy metabolism showed evidence of artificial selection during the domestication of laboratory strains. Our findings demystify the puzzling origin and migration of brown rats and reveal the impact of evolution and domestication on this animal.

## Introduction

A detailed understanding of the geographic origin of the wild rodents and its subsequent dispersal routes across the globe would be informative in clarifying the spread of diseases and human migration (Matisoo-Smith and Robins 2004; Lin et al.2012). For example, study from mtDNA phylogenies of the Pacific rat (*Rattus exulans*) revealed origins and dispersals of Pacific peoples (Matisoo-Smith and Robins 2004). Study of mtDNA indicated that the house mice (*Mus musculus domesticus*) were a valuable proxy for Viking movements (Searle et al. 2009). Phylogeographical analysis on the black rats (*Rattus rattus*) in the western Indian Ocean revealed that the history of black rats was compatible with human colonization history (Tollenaere et al. 2010).

The brown rat (*Rattus norvegicus*), one of the most common rats whose habitat overlaps with that of human, is undesirable in human history, acting as a reservoir for a number of zoonotic pathogens such as Hantavirus and disseminating many diseases (Meerburg et al. 2009; Lin et al. 2012; Kosoy et al. 2015). It is well-recognized that the brown rat spread out of Asia to Europe (Silver 1941; Southern 1964; Freye et al. 1968; Amori and Cristaldi 1999; Kosoy et al. 2015). This have been supported by archaeological data (Suckow et al. 2006) and genetic evidences from both mitochondrial and nuclear markers (Song et al. 2014; Puckett et al. 2016). However, until now, the detailed geographic origin and the dispersal route of brown rats from Asia to Europe have remained controversial. Based on the evidences of fossils and bones, it was suggested that *Rattus norvegicus* originated in Northeast China and Southeast Siberia (Wilson and Reeder 2005; Ness et al. 2012; Kosoy et al. 2015), and then dispersed westward through the Eurasian steppes into Europe (Gibbs et al. 2004). However, a recent study, based on part of mtDNA sequences, alludes to southern China as a possible origin of the brown rat migration (Song et al. 2014), creating the possibility that it might have dispersed alongside humans from southern East Asia to other regions.

Likewise, when brown rats arrived at Europe/Africa remains controversial. Historical records, based mainly on rat bones, indicated that brown rats appeared in Europe during the Medieval Ages and became widespread during the Industrial Revolution (Amori and Cristaldi 1999). Suckow et al (2006) suggested arrival dates for the rats in Ireland, England, France, Germany, and Spain of, 1722, 1730, 1735, 1750, and 1800AD, respectively. However, evidence also suggested that brown rats might have been present in Europe as early as 1553AD (Freye et al. 1968), and were introduced into North America by the 1750s (Armitage 1993).

Studies using mitochondrial DNA or a limited number of nuclear markers might be biased in inferring the geographical origins and migratory routes of human and animals. Recently, the rapid development of next generation sequencing and population genetics, has facilitated studies to clarify the origin and migration of humans as well as domestic animals at a genome wide scale (Pagani et al. 2015; Frantz et al. Malaspinas et al. 2016; Mallick et al. 2016; Wang et al. 2016). To clarify the origin and migration of brown rats and selection on their genomes, in this study we sequenced the whole genomes of 110 rats across the world, including China (N=38), Russia (N=5), Southeast Asia (N=8), Middle East (N=12), Europe (N=26) and Africa (N=21), as well as the genomes of 8 outgroup rats. Based on these genomes, we showed that brown rats migrated about 3600 years ago from southern East Asia, rather than Northern Asia as formerly suggested, to the Middle East and then to Europe and Africa. Our study provides a model of “out of southern East Asia” for the brown rats, which shifts our perspectives on the origin and migration of brown rats.

The laboratory rat, domesticated from the wild brown rat and widely used in biomedical research as an animal model for ~160 years, is commonly believed to have been first used in experiments in Europe in 1856 (Suckow et al. 2006). Compared to their wild ancestors, laboratory rats exhibit remarkable differences in their morphological, physiological and behavioral attributes such as coat color, organ size, energy metabolism, reproductive performance and tameness (Whishaw and Kolb 2004; Baker et al. 2013). However, the genetic mechanisms underlying these phenotypic differences are still unclear. By comparing genomic and transcriptomic data between wild brown rats and laboratory rats (Atanur et al, 2013), we found that many genes involved in the nervous system (e.g. *EGR2, PTGDS, CLOCK* and *FOXP2*) and energy metabolism (e.g. *ATP5D, COX7A2, COX8A, NDUFB2* and mitochondrial genes) in the laboratory rat have adaptively evolved and exhibit differential expression, indicating that these candidate genes are likely associated with changes in the behavior of the laboratory rats related to domestication, and might have facilitated the successful domestication of the brown rats.

## Results and Discussion

### Out of southern East Asia of wild brown rats

To understand the origin and dispersal routes of the brown rat, a total of 117 *Rattus norvegicus* samples across the world and 1 black rat (*R. rattus*) were collected for genome sequencing. The species status of these *Rattus norvegicus* was carefully identified by morphology and further confirmed with cytochrome *b* (*cytb*) sequences by Sanger sequencing. However, based on whole genome sequence data, the specific status of 7 individuals was ambiguous as they showed a closer relationship with the black rat (*R. rattus*). These seven individuals, along with the black rat were treated as the outgroup for the following analyses. Whole genome sequences were therefore generated for 110 brown rats from China (N=38), Russia (N=5), Southeast Asia (N=8), Middle East (N=12), Europe (N=26) and Africa (N=21), as well as 8 outgroup rats (**Supplemental Fig. S1, Supplemental Table S1**).

The genetic diversity of the wild rats varied with geography, with rats from Asia harboring the highest levels of diversity, supporting an out-of-Asia origin (**Supplemental Table S2**). To identify the primary source and dispersal time of the wild rats, we performed analyses that included phylogenetic construction (**Fig. 1A**, **Supplemental Fig. S2-S6**), principal components (**Fig. 1B**), Bayesian clustering (**Supplemental Fig. S7**), haplotype sharing analyses (**Supplemental Fig. S8**) and demographic modelling (**Supplemental Fig. S9-S19, Supplementary Note**). The phylogenetic tree, capable of investigating hypotheses of evolutionary history, was constructed by the neighbor-joining method using 24,977,888 autosomal single nucleotide polymorphisms (SNPs) identified in our genome data (**Fig. 1A**, **Supplemental Fig. S4A**). Rats from outside Asia clearly showed closer relationships with rats from southern East Asia (including Southeast Asia and southern China), than to those from Northern Asia (including northern China and Russia). The maximum likelihood tree constructed using TreeMix (Pickrell and Pritchard 2012) from the SNPs data supported the above phylogeny (**Supplemental Fig. S20**). We also employed approaches implemented in mega, raxml, fasttree and rapidNJ based on the Maximum Likelihood (ML) or Neighbor-Joining methods to construct phylogenetic trees (Simonsen et al. 2008; Price et al. 2010; Stamatakis 2014; Kumar et al. 2016) (**Supplemental Fig. S2-S6**), and obtained similar results, with rats from outside Asia clearly showing closer relationships with rats from southern East Asia than to those from Northern Asia.

**Figure 1.**
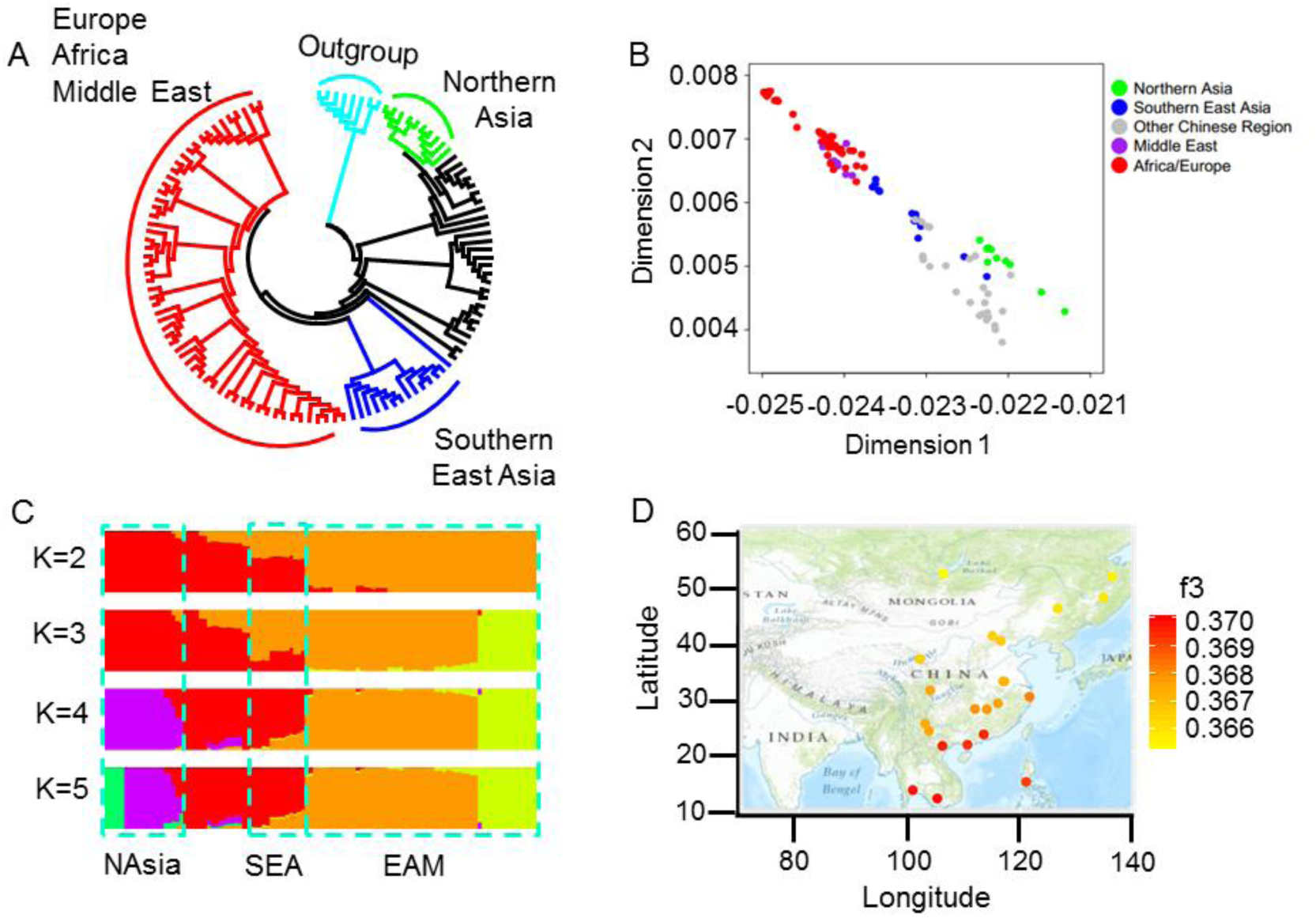
Out of southern East Asia origin of wild brown rats. (A) Phylogenetic neighbor-joining tree. Black colored lines represent samples from China. (B) PCA analysis. (C) Bayesian clustering analysis by ADMIXTURE. NAsia (northern Asia), SEA (southern East Asia), EAM (Europe, Africa, Middle East). (D) f3 -outgroup statistics showing genetic proximity of Europe population to Eastern Asia individuals.

Principal components analysis (PCA) positioned the southern East Asian rats between the European/African/Middle East and Northern Asian rats (**Fig. 1B**), corroborating the clustering of the European/African/Middle East rats with the southern East Asian rats seen in the neighbor-joining tree (**Fig. 1A**). This pattern was again supported by a Bayesian clustering analysis (Alexander et al. 2009), which further indicated that the European/African/Middle East rats have a more recent common ancestry with populations from southern East Asia, rather with those from Northern Asia (**Fig. 1C**, **Supplemental Fig. S7**). The f3-outgroup statistic showed that European/African/Middle Eastern rats harbored a clear genetic similarity with southern East Asian rats (**Fig. 1D**, **Supplemental Fig. S21**). This was further confirmed by a haplotype sharing analysis, where the Out-of-Asia rats showed more proximity to southern China rats than those from northern China (**Supplemental Fig. S8**).

In contrast to the hypothesis that the wild brown rat dispersed from northern Asia to Europe (Gibbs et al. 2004), all of our analyses support the alternative hypothesis that wild brown rats dispersed from southern East Asia to Europe/Africa/Middle East.

To further clarify whether Northern Asia or southern East Asia was the birthplace of wild brown rats, we simulated demographic models based on maximum likelihood method (Li and Stephan 2006; Excoffier et al. 2013), and found that southern East Asia was more likely to be the cradle, namely, brown rats migrated from southern East Asia to Northern Asia (**Supplementary Note, Supplemental Fig. S11-S12, Supplemental Table S3-5**), consistent with previous study (Song et al. 2014). It should be noted that although the northern Asia population is closer to the outgroup than the southern East Asia population in the phylogenetic tree (**Fig. 1A**), we cannot make the conclusion of the northern Asia as the birthplace of wild brown rats because the two migrations out of southern East Asia can cause the same tree branching pattern. Given the demographic model of southern East Asia as the birthplace, the simulated phylogenetic trees (representing the true evolutionary history) were always rooted at the branch of the northern Asia population (**Supplemental Fig. S6**).

### Dating two migrations out of southern East Asia

We then dated the migration of brown rats out of southern East Asia using the well-established maximum likelihood method based on the joint site frequency spectrum (Li and Stephan 2006; Excoffier et al. 2013). We found that brown rats migrated from southern East Asia to Northern Asia about 201,800 years ago, while brown rats spread from southern East Asia to the Middle East ~3600 years ago (95% CI: 3100-5800), to Africa ~2600 (2100-3300) years ago and to Europe ~1800 (1200-2600) years ago (**Fig. 2**, **Supplemental Note, Supplemental Fig. S9-S19, Supplemental Table S2, Supplemental Table S6-S12**). We also performed a robustness analysis by re-estimating the migration times using different generation times (2 generations per year and 3 generations per year) (Anderson 1967)for the rat and different divergence times between rat and mouse (**Supplementary Note**). The re-estimated introduction times vary but generally fall into the confidence intervals obtained above (**Supplemental Fig. S18, and Supplemental Table S13-S14).**

**Figure 2.**
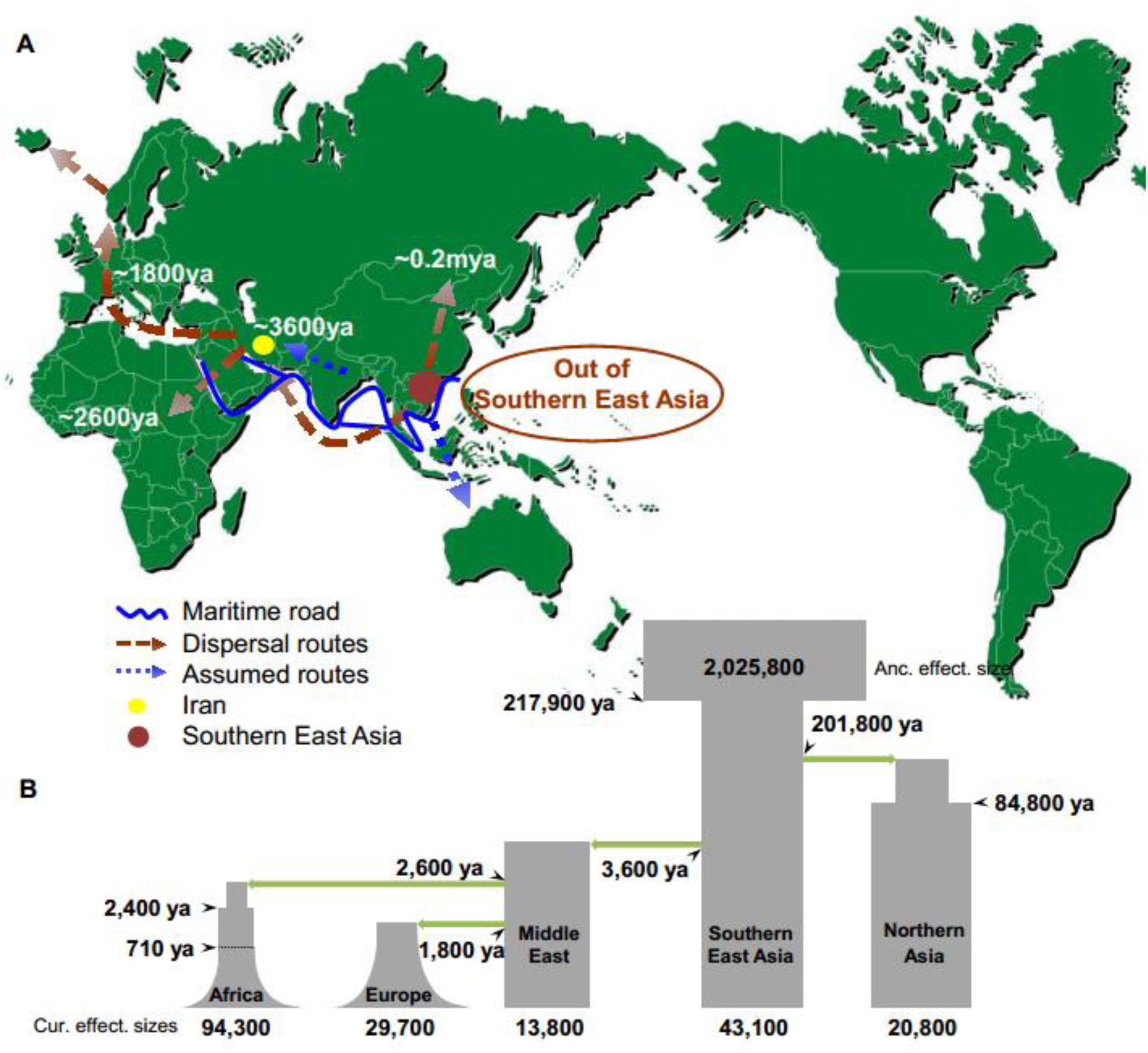
Dispersal routes and demographic histories of wild brown rats. (A)Proposed dispersal routes of the wild brown rats based on our analyses. (B) The inferred joint demographic model of different wild brown rats populations based on the maximum likelihood method.

The estimates of the introduction times of brown rats to Europe are much older than historical reports that propose migrations in the 18th century. However, we shall note that the maritime trade has been in existence in the Indian Ocean and southern East Asia region for over 4,000 years (Forbes 1995; Miksic 2013), and those early human activities could have facilitated the migration and dispersal of brown rat from southern Asia to other regions. Such kind of human assisted migration/introduction was often proposed for rodents. For example, the pacific rat (*R. exulans*) and the black rat (*R. rattus*), migrated out of southern East Asia to remote areas of Oceania and Madagascar, respectively, more than 3000 years ago (Matisoo-Smith and Robins 2004; Tollenaere et al. 2010), supporting the association between human navigation and the migration of rats.

A caveat of this analysis is the small population sizes and low depth of some genome sequences used in some of our analyses. Further studies with larger population sizes and more populations (with high depth and coverage genomes) should undoubtedly help to refine the dispersal times for the brown rats. In addition, to reconcile the genetic and archaeological evidences, archaeological studies are needed to focus on more ancient bones. Genome sequencing of ancient bones of brown rats from Europe, Africa and Middle Asia could also be employed to clarify the dispersal routes and times.

### Rapid evolution of immune response genes in wild brown rat population

During dispersal, wild rats have transmitted and spread devastating diseases to human populations. This property of rats, allowing them to host many pathogens, has long remained a puzzle. An “arms-race” that drives the rapid evolution of the immune system in a host (Van Valen 1973) might have endowed rats with this potential. Therefore, we investigated whether genes involved in the immune system might have rapidly evolved, potentially under positive selection, in wild brown rats during their dispersal. As expected, when we examined differentiation of genes between European and Chinese wild brown rats, a large number of genes with significantly high level of population differentiation were enriched for functions related to the immune system, such as “*leukocyte mediated immunity*”, “*response to bacterium^1^* and “*leukocyte mediated cytotoxicity*” (**Fig. 3**, **Supplemental Table S15).** A comparison of African and Chinese rat genomes yielded a similar finding (**Supplemental Table S16**). In particular, the gene *Mgat5* showed the highest level of population differentiation between the Chinese and European rats (**Fig. 3**). *Mgat5* participated in the synthesis of galectins, cell-surface ligands involved in T-cell proliferation. *Mgat5*-knock-out mice display an autoimmune phenotype, and loss of *Mgat5* lowers the threshold needed for activation of T-cells (Demetriou et al. 2001). The window with the second highest level of differentiation contained single gene, *Lyst*, whose mutations causes the Chediak-Higashi Syndrome in human, a genetic immunodeficiency disease where T-cell and natural killer cell cytotoxicity become defective (Trantow et al. 2010). The differentiation of immune genes suggests that wild rats from different regions of the world might differ in their susceptibility to specific pathogens, a hypothesis that needs experimental validation.

**Figure 3.**
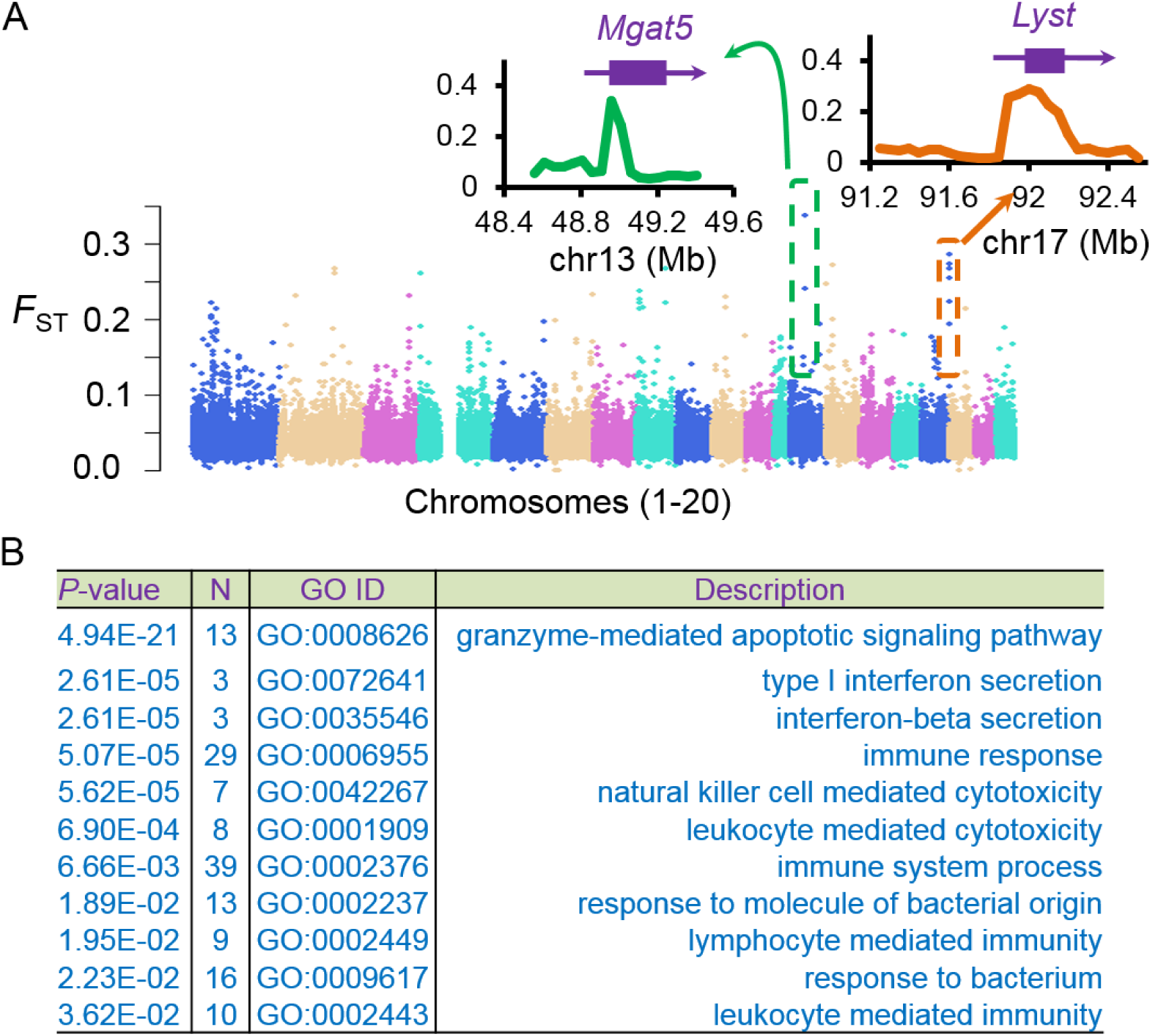
Immune response genes are under selection in the wild brown rats. (A)Genomic landscape of the population differentiation (*F*_st_) between European and Chinese rats. Top two clusters with high level of population differentiation across genes *Mgat5* and *Lyst* were represented. (B) List of genes with significantly high level of population differentiation between European and Chinese rats are enriched for immune system related functions. Gene Ontology analysis of the protein-coding genes was conducted using an online annotation tool g:Profiler and P-values were corrected by Benjamini-Hochberg FDR (Reimand et al. 2011).

### Artificial selection during the domestication of laboratory rat

The laboratory rat, domesticated from the wild brown rat, has been widely used in biomedical research as an animal model for ~160 years, since it was believed to be first used for experiment in Europe in 1856 (Suckow et al. 2006). However, specific origin of the laboratory rat and the genetic relationship between inbred strains and wild rats has not been clearly determined. Our analyses, including phylogenetic tree construction (**Supplemental Fig. S22**), ADMIXTURE (**Supplemental Fig. S23**), PCA (**Supplemental Fig. S24**), and f3-outgroup statistic (**Supplemental Fig. S25**), support a single origin for the laboratory rat (Kuramoto et al. 2012), from a sister group to the Europe/Africa/Middle East rat group. Laboratory rats were domesticated from wild rats and may experience a strong bottleneck. Therefore, it is difficult to narrow down the specific origin of laboratory rats based on our genome data.

Compared to their wild ancestors, laboratory rats exhibit remarkable differences in their morphological, physiological and behavioral attributes such as coat color, organ size, reproductive performance and tameness (Whishaw and Kolb 2004; Baker et al. 2013). It makes the laboratory rat as an ideal model to study the mechanism of domestication.

We evaluated the differentiation of each SNP and performed a sliding window analysis to identify regions/genes harboring high levels of differentiation between laboratory and wild rats. A total of 292 candidate genes (**Supplemental Table S17-19**) were identify to show *F*_ST_ values in the top 1%, and thus were considered to be potentially positively selected. Many of these genes were involved in sensory perception, consistent with a differing ability in sensory perception. In addition, a gene enrichment analysis found that many of the genes have function in “*neurological system process*” (54 genes, GO: 0050877, P=2.11×10^−5^ corrected by Benjamini-Hochberg FDR, **Supplemental Table S19**). In particular, seven of the genes (*AFF2, MECP2, NAA10, NSDHL, SLC6A8, SLITRK1, ENSRNOG00000049488*) were enriched at the HPO category of “*abnormally aggressive, impulsive or violent behavior*” (P=0.05, corrected by Benjamini-Hochberg FDR, **Supplemental Table S19**), which might explain the behavioral changes observed in laboratory compared to wild rats. As observed in other domesticated animals (Li et al. 2013; Carneiro et al. 2014), positive selection on genes involved in the nervous system might have played key roles in the successful domestication of the laboratory rat from wild brown rat ancestor. When the expression levels of selected candidate genes was examined it was found that the expression level of the positively selected genes, *CLOCK*, a central regulator in the regulation of circadian rhythms (Vitaterna et al. 1994), and *FOXP2*, a central gene in vocal behavior (Enard et al. 2002), were significantly up-regulated in the hypothalamus of the laboratory rats compared to the wild brown rats (**Supplemental Fig. S26**), concordant with the change in circadian rhythms and behavior.

Genomic loci under artificial selection possess other distinctive patterns such as low genetic diversity and long haplotype homozygosity (Sabeti et al. 2006). We therefore sought signals of artificial selection in the laboratory rats using cross-population extended haplotype homozygosity (XP-EHH). In total, 447 protein coding genes were identified as candidate positively selected genes (**Supplemental Table S20**) which showed high XP-EHH values that were in the top 1%. Gene enrichment analysis found that 14 of these genes are enriched in the category of “*regulation of developmental growth*” (GO:0048638, P=0.002, **Supplemental Table S21**). Selection on these developmental genes might be associated with many of the morphological differences observed between laboratory and wild rats. For example, laboratory rats are larger and have weaker bone structure and smaller internal organs (including brain, heart, liver, and spleen) (Stryjek et al. 2012). The top 5 windows showing the highest XP-EHH values clustered together and overlapped with the protein-coding gene, *B3GAT1* (**Fig. 4A**, P=1.96×10^−5^). *B3GAT1* is involved in the biosynthesis of HNK1 (Mitsumoto et al. 2000), which is widely expressed in the brain, and knockout mice exhibit reduced long-term potentiation at Schaffer collateral-CA1 synapses and defects in spatial learning and memory (Yamamoto et al. 2002). The level of mRNA expression of *B3GAT1* was significantly up-regulated in the brain of laboratory rats compared to wild brown rats (**Fig. 4A**). We propose that the up-regulation of *B3GAT1* likely enhanced spatial learning and memory in laboratory rats, enabling them to adapt to captivity as part of the process of domestication. Generally, wild animals are more active and reactive and show extreme levels of stress in captive environments, properties that contribute to their higher mortality in captivity (Price 2002). Increased spatial learning and memory, due to changes in *B3GAT1*, might have helped to reduce stress levels in domesticated animals.

**Figure 4.**
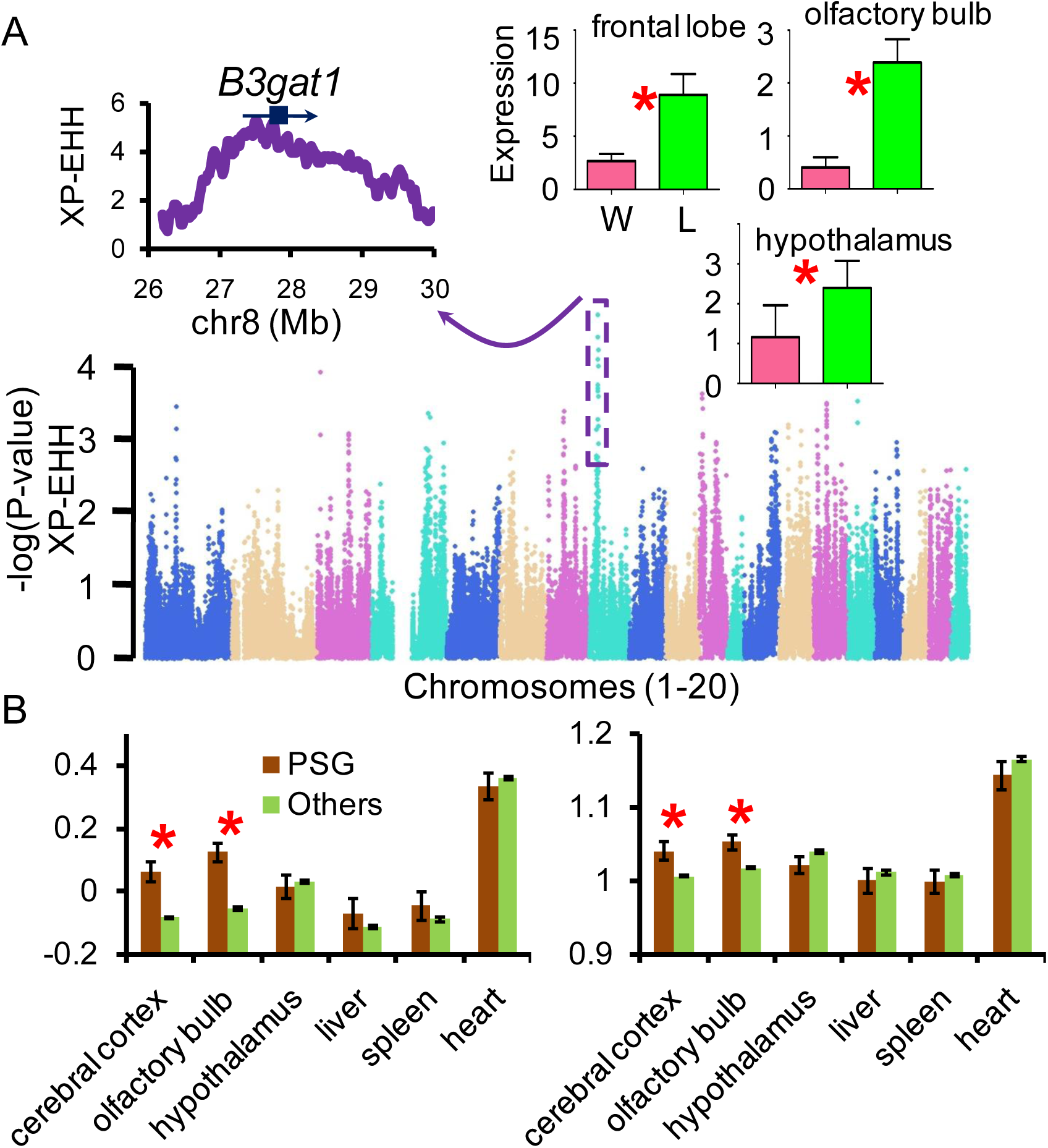
Artificial selection in laboratory rats. (A) Genomic landscape of the -log10(p-value) of XP-EHH values of laboratory rats compared with wild brown rats and selection at the *B3GAT1* gene. Increased mRNA expression levels of *B3GAT1* gene detected by RT-PCR in the frontal lobe, olfactory bulb and hypothalamus of laboratory rats (L) compared to wild brown rats (W). Asterisk identifies statistically significant differences (P<0.05). (B) Comparisons of the difference in the levels of gene mRNA expression between wild brown and laboratory rats. Y axis represents the mean expression difference (±S.E.) of positively selected genes (PSG) and other genes. The expression value for each gene was transformed by (log2(FPKM+1)). Left: Expression difference of each gene calculated as the difference between wild brown rat and laboratory rat values. Right: Difference of each gene was calculated by (Ewild + 1)/(ELaboratory +1).

Changes in gene expression play very important roles in phenotypic evolution. We proposed that genes displaying differential expression between laboratory and wild brown rats might account for phenotypic changes necessary for the domestication of the laboratory rat. To better understand the genetic basis of these phenotypic differences, transcriptomes of the cerebral cortex, hypothalamus, olfactory bulb, liver, spleen and heart were profiled by RNA-sequencing from wild brown and laboratory rats. Genes that showed high levels of population differentiation between wild brown and laboratory rats also presented with significantly higher levels of differential gene expression in the cerebral cortex and the olfactory bulb (*P*= 5.44×10^−7^, 6.42×10^−7^, **Fig. 4B (left)**, *P*=8.31×10^−7^, 3.38×10^−6^, **Fig. 4B (right)**, Mann-Whitney U test). However, significant differences in expression patterns were not found for other tissues **(Fig. 4B)**. Differential expression of these differentiated genes in the cerebral cortex and the olfactory bulb provided a plausible explanation for changes in behavior and sensory abilities of laboratory rats compared to wild brown rats.

Overall, 97 genes were found to be differentially expressed in the cerebral cortex between wild brown and laboratory rats, a number much higher than any other tissue (**Supplemental Fig. S27**). In a gene enrichment analysis, 10 of these 97 genes fell in the category “*behavior*” (GO: 0007610, *P*= 3.19×10^−2^, **Supplemental Table S22-S23**). Differentially expressed genes were also significantly over-represented in KEGG categories related to brain disorders, such as “*Alzheimer’s disease*”, “*Parkinson’s disease*”, and “*Huntington’s disease*”. Changes in the expression levels of these genes might have facilitated the domestication of laboratory rats by influencing the evolution of the nervous system, as seen in other domestic animals, such as dogs and rabbits (Li et al. 2013; Carneiro et al. 2014).

### Differential expression of genes involved in energy metabolism in the brain between wild brown and laboratory rats

A notable feature of the transcriptome data was that many of the differentially expressed genes associated with energy metabolism were over-represented in categories like “*oxidative phosphorylation*”, “*respiratory electron transport chain*”, “*oxidation-reduction process*”, and “*generation of precursor metabolites and energy*” (**Supplemental Table S22-S23**). Assessing changes in energy metabolism and their consequent nervous system disorders is a key pillar in evolutionary studies of the nervous system (e.g. human brain). Genes related to energy metabolism have been implicated in both the evolution and maintenance of human-specific cognitive abilities (Khaitovich et al. 2008). Since the mitochondrion is the energy producing machinery of a cell, we examined the expression of 11 mitochondrial protein-coding genes and found they harbored significant differential expression levels in frontal lobe tissue between the laboratory and wild brown rat (**Fig. 5A**). Except for *MT-ND3*, all other identified mtDNA genes were upregulated in the laboratory rat; however, no significant difference in the copy number of mitochondrial DNA was found between the laboratory and wild brown rats (**Fig. 5B**). In contrast, only *ND3* demonstrated significant differential expression in heart and liver tissues. These data suggest that substantial evolutionary change in energy metabolism has occurred in the brain of the laboratory rat since domestication.

**Figure 5.**
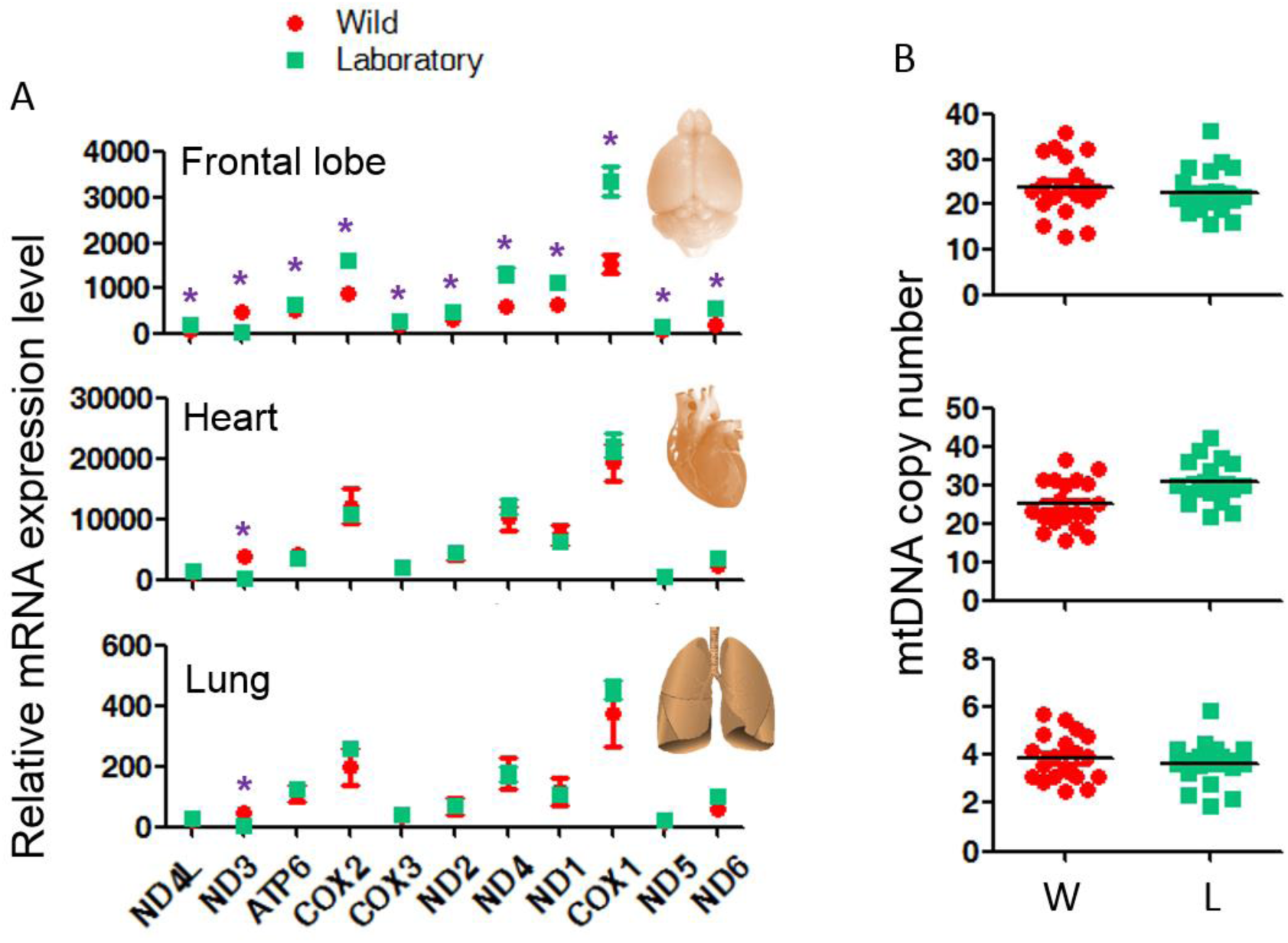
mRNA expression levels of mitochondrial coding genes (A) and mtDNA copy number (B) in wild brown (W, red) and laboratory (L, green) rats. Asterisk (*) indicates that expression level is statistically significant differential expression (p<0.05).

## Conclusion

In conclusion, we provide evidence for an out of southern East Asia origin for the brown rat and its subsequent dispersals to the Middle East, Europe and Africa thousands of years ago. Along with the migration, many genes involved in immune response have adaptively evolved under natural selection in the wild rats. Compared to wild brown rats, many genes involved in the nervous system and energy metabolism have evolved selectively and exhibit differential expression in the laboratory rat, suggesting that these genes are associated with the changes in behavior of the laboratory rats associated with domestication.

## Methods

The handling of animals used in this study followed the guidelines and regulations of the Kunming Institute of Zoology on animal experimentation and was approved by the Institutional Animal Care and Use Committee of the Kunming Institute of Zoology.

### DNA samples for genome sequencing

A total of 117 (putative) *Rattus norvegicus* samples from Russia, Northern China, Southern China, Southeast Asia, Europe, Africa, and Middle East, and 1 black rat (*R. rattus*) were collected for genome sequencing. The species status of the *Rattus norvegicus* samples were assessed by morphology and by Sanger sequencing of cytochrome *b* (*cytb*) sequences.

Whole genome sequences of 24 laboratory rats except Brown Norway breed were obtained from a previously published study (Atanur et al. 2013) to discern potential artificial selection in the genome of laboratory rats. As the genomes of the laboratory rats were completed at a higher depth (>20X), the genomes of 6 our wild brown rats, from Mali (N=1), Morocco (N=1), Russia (N=2) and China (N=2), were sequenced at high depth (~20X) for the comparison with the laboratory rats.

### Genome sequencing

Tissues for DNA extraction were stored in alcohol at -80°C. Ten micrograms of genomic DNA, prepared by the standard phenol chloroform extraction protocol, was used to construct libraries with a 350 base pair insert size. Sequencing libraries were generated using the NEB Next^®^ Ultra DNA Library Prep Kit for Illumina^®^ (NEB, USA) following manufacturer’s recommendations and index codes were added to attribute sequences to each sample. Briefly, Chip DNA was purified using the AMPure XP system (Beckman Coulter, Beverly, USA). After adenylation of the 3’ ends of the DNA fragments, NEB Next Adaptors with hairpin loop structure were ligated to prepare for hybridization. Electrophoresis was then used to select DNA fragments of a specified length. 3 μL USER Enzyme (NEB, USA) was used with the size-selected, adaptor-ligated DNA at 37°C for 15 min followed by 5 min at 95°C before PCR. PCR was performed with Phusion High-Fidelity DNA polymerase, Universal PCR primers and Index (X) Primer. Finally, PCR products were purified (AMPure XP system) and library quality was assessed on the Agilent Bioanalyzer 2100 system. Clustering of index-coded samples was performed on a cBot Cluster Generation System using HiSeq 2500 PE Cluster Kit (Illumia) according to the manufacturer’s instructions. After cluster generation, library preparations were sequenced on an Illumina Hiseq 2500 platform and 125 bp paired-end reads were generated. Raw data from sequencing contains many low quality reads with adapters. We filtered these reads to obtain clean reads as follows:

1. Remove read pairs containing adapter;
2. Remove read pairs generating a single sequence where the N content is greater than 10% of the read length;
3. Remove read pairs generating a single sequence where the number of low quality of base (Q<5) is greater than half of the reads.

### Read mapping, SNPs calling, filtering and imputation

Quality filtered reads were mapped to the reference *Rattus norvegicus* genome rn5 (ENSEMBL version 72) (Gibbs et al. 2004) using the alignment algorithm BWA-MEM (Li 2013). Single nucleotide polymorphisms were detected using the Genome Analysis Toolkit (McKenna et al. 2010). Duplicate read pairs were first identified using the Picard tools (http://picard.sourceforge.net/), then realigned around putative indels, which were downloaded from dbSNP. To provide empirically accurate base quality scores for each base in the read pairs, we performed base quality recalibration and to reduce false positive rate, we applied hard filters according to GATK guidance. Here are the criteria we used to filter the raw SNPs: QD < 2.0, FS > 60.0, MQ < 40.0, HaplotypeScore > 13.0, MappingQualityRankSum < −12.5, ReadPosRankSum < −8.0, and -cluster 3 -window 10. As the genome sequences used in our dataset were of low depth, we performed imputation of ungenotyped markers using BEAGLE (Browning and Browning 2009) after removing triallelic sites and GATK filtering of the raw vcf file. Finally, a total of 24,977,888 autosomal SNPs were obtained.

### Phylogenetic relationships and population structure analysis

Phylogenetic relationships using the complete SNPs dataset were constructed by the neighbor joining method using the rapidNJ method (Simonsen et al. 2008). Considering computer memory consumption and the effects of LD, we also constructed phylogenetic trees with bootstrap support values (**Supplemental Fig. S2-S5.**) after thinning the SNP data sets at a 50k (using vcftools with –thin 50000 parameter and 49,196 SNPs left),10k (using vcftools with –thin 10000 parameter and 225,012 SNPs left) and 1k (using vcftools with –thin 1000 parameter and 1,628,064 SNPs left) distance using the neighbor-joining or maximum likelihood methods using MEGA, raxml, fasttree, **TREEMIX** and rapidNJ (Simonsen et al. 2008; Price et al. 2010; Pickrell and Pritchard 2012; Stamatakis 2014; Kumar et al. 2016) with the different SNP datasets.

To reveal the relationships among different geographical locations, we performed a principle components analysis (PCA) using GCTA (Yang et al. 2011) with the complete SNPs dataset after PLINK (Purcell et al. 2007) format conversion. Subsequently, population structure was deduced by the program ADMIXTURE, a tool for maximum likelihood estimation of individual ancestries from multi locus SNP genotype datasets (Alexander et al. 2009), with different K values (from 2 to 10) and –cv -B flags. A good value of K (K=5) exhibited a low cross-validation (CV) error compared to other K values, and -B flag is a basic way to obtain bootstrap standard errors. Analysis of haplotype sharing between the different populations was performed as described in vonHoldt et al. (2010) with non-overlapping 20k windows. If there were more than 5 SNPs in a window, then we randomly selected 5 of these SNPs.

### Outgroup f3 analysis

To further assess the genetic proximity of non-Asian populations to Asian individuals, we computed the outgroup f3-statistic (Patterson et al. 2012) to estimate their shared genetic history, and identify individuals showing a closer relationship with the Southern East Asian individuals.

### Analysis of the signatures of positive selection

The population pairwise estimate of differentiation (*F*_ST_) was calculated as previously described (Akey et al. 2002). Sliding window analysis was performed with a window size of 100kb, and a step size of 50kb. For the analysis of extended haplotype homozygosity, haplotypes of each chromosome were deduced using the software SHAPEIT (Delaneau et al. 2013) and XP-EHH values were calculated with the software XPEHH (Sabeti et al. 2007) (http://hgdp.uchicago.edu/Software/). Sliding window analysis was performed with a window size of 100kb, and a step size of 50kb.

### RNA sequencing and analysis

Tissues for RNA-seq, including heart, liver, spleen, cerebral cortex, hypothalamus and olfactory bulb, from a Sprague dawley (SD) rat and a wild brown rat were stored in RNAlater (Ambion, Austin, TX, USA). Total RNA was extracted using the standard Trizol (Qiagen, Chatsworth, CA, USA) protocol and RNeasy mini kit (Qiagen, Chatsworth, CA, USA). Before library construction, we assessed the quality of the RNA by spectrophotometry using NanoDrop 2000, gel electrophoresis and Agilent 2100. The library was prepared following the Illumina Genomic RNA sample prep kit protocol and then sequenced on an Illumina HiSeq 2000 platform following the manufacturer’s instructions.

Adapter sequences were first removed from our own RNA-seq raw data using Cutadapt (v1.2.1) (Martin 2011). Before alignment, reads were trimmed based on their quality scores using the quality trimming program Btrim (Kong 2011). Reads were aligned to the rat reference genome (rn5) (Gibbs et al. 2004) using TopHat (v2.0.4) (Trapnell et al. 2009) and then assembled using Cufflinks (v2.0.2 with -g parameter) (Trapnell et al. 2012). The differential expression of genes in the different tissues was calculated using Cuffdiff (Trapnell et al. 2012).

To assess if difference in the levels of gene expression in the heart, liver, spleen, cerebral cortex, hypothalamus and olfactory bulb observed between wild brown and laboratory rat might have been driven by positive selection at local regulatory sites during domestication, a series of statistical tests were performed. Expression levels (FPKM) for each gene in each tissue were retrieved and transformed as log2(FPKM+1). Differences in expression levels for each gene between the wild brown rat and the laboratory rat were calculated using log2((FPKMwild +1) /(FPKMlab +1)). We then compared the differences in the expression levels of positively selected genes (PSGs), identified based on their significant *F*_ST_ and XP-EHH values, to all other genes (all genes in the whole genome excluding PSGs) by the Mann-Whitney U test.

### Real-time quantitative PCR (qPCR) of selected genes and energy metabolism related genes

We synthesized single-stranded cDNA from 1 μg of total RNA using the PrimeScript RT-PCR Kit (TaKaRa, Japan) in a 25 μL final reaction volume according to the manufacturer’s instructions. To validate the differential expression of genes detected in the above RNA-seq analysis, the relative abundance of the mRNAs for eight genes involved in the nervous system and energy metabolism, i.e. *ATP5D, CLOCK, COX7A2, COX8A, EGR2, FOXP2, NDUFB2* and *PTGDS* genes were measured by the qPCR using samples of RNA from the cerebral cortex, olfactory bulb and hypothalamus from three wild brown and three laboratory rats. The relative standard curve method was applied with normalization to the housekeeping gene β-actin.

### Detection of mRNA expression of mitochondrial protein coding genes and mitochondrial DNA copy number

The relative abundance of the mRNAs encoding 11 mitochondrial protein-coding genes was estimated in the cerebral cortex, heart and lung from three wild brown and three laboratory rats by qPCR as described above. To detect mitochondrial DNA copy number, genomic DNA was extracted by the Genomic DNA Miniprep Kit (Axygen, AP-MN-BL-GDNA-250), and mitochondrial DNA copy number was measured using real-time quantitative PCR with samples of cerebral cortex, heart and lung from ten wild brown and ten laboratory rats, with normalization to the *Hbb* (β-globin) gene. qPCR was performed on the iQ2 system platform (BioRad Laboratories) with SYBR Premix Ex Taq II kit (TaKaRa, DRR081A).

### Analysis of functional term enrichment

Gene Ontology analysis of the protein-coding genes was conducted using an online annotation tool g:Profiler and P-values were corrected by Benjamini-Hochberg FDR (Reimand et al. 2011), which used genome wide genes as background. Expressed genes were also used as background gene set in each tissue for enrichment analysis of differentially expressed genes by web-based DAVID (Dennis et al. 2003).

### Substitution rate and the robustness analysis for generation time and divergence time between rat and mouse

Since substitution rate is generally faster in the rodent lineage than many other mammals (Wu and Li 1985), we estimated substitution rate based on the pairwise genome alignment between the rat (*Rattus norvegicus*, rn5) and the house mouse (*Mus musculus*, mm10). A total of 1,720,780,766 sites were in the alignment (indels and sites containing ‘N’ (ambiguous nucleotide) were not taken into account) with 257,482,102 substitutions observed since rat and house mouse divergence. The mean divergence time between mouse and rat is about 22.6 million years ago according to TIMETREE (Hedges et al. 2006; Kumar and Hedges 2011; Hedges et al. 2015). If we assume the rat has 2 generations per year (Ness et al. 2012; Halligan et al. 2013; Deinum et al. 2015), we estimate a genome-wide nucleotide substitution rate is 1.655×10^−9^ per generation per base pair. If three generations per year is assumed (Ness et al. 2012; Halligan et al. 2013; Deinum et al. 2015), the estimated substitution is reduced to 1.103×10^−9^, and this rate is also used to examine the robustness of our estimated introduction times. Based on 47 studies of nuclear genes in TIMETREE (Hedges et al. 2006; Kumar and Hedges 2011; Hedges et al. 2015), the estimated divergence time between mouse and rat varies (**Supplemental Table S14**). Thus we re-estimated introduction times based on a range of mouse-rat divergence times range (we used a range of divergence times from 15 to 30mya).

### Maximum likelihood inference of the demographic history of brown rats

To obtain the joint site frequency spectrum (SFS) (Li and Stephan 2006; Gutenkunst et al. 2009), the ancestral state of each allele was inferred using the reference genome of the house mouse (mm10). Here, we only considered bi-allelic sites, and indels were excluded. The unfolded SFS was also obtained for each population.

The previously proposed likelihood function for demographic history (Li and Stephan 2006; Excoffier et al. 2013) was calculated using fastsimcoal2 (Excoffier et al. 2013). The ranges of parameters in the configure-file for fastsimcoal2 are listed in **Supplemental Table S12**. For each demographic scenario, 100,000 simulations per likelihood estimation (–n 100000 -N 100000), at least 20 ECM (expectation-conditional maximization) cycles (-l20), at most 40 ECM cycles (-L40) were used as the command line parameters in each run. To avoid being trapped in a local optimum, 400~2,000 runs were applied. The Akaike information criterion (AIC) information criterion was used to compare the different models, with AIC=2k-2ln (MaxEstLhood), where *k* is the number of parameters to be estimated in each model, and MaxEstLhood is the maximum value of the likelihood function for each model.

To obtain confidence intervals of the estimates, 100 independent DNA polymorphism datasets (as joint SFS) were simulated conditional on the estimated demographic parameters. The maximum likelihood analysis was then applied to each joint SFS with 30 ECM cycles and 30 runs, and 100,000 coalescent simulations were conducted to calculate the likelihoods. The empirical distributions of the estimates and the 95% confidence intervals were then obtained.

To further examine how the real data fit the chosen final joint demographic model, we followed methods developed by others (Excoffier et al. 2013) and applied an approach based on a likelihood ratio G-statistic (Nielsen et al. 2005; Nielsen et al. 2009). The composite likelihood ratio (CLR) is calculated as CLR=log10 (CLo/CLe), where CLo is the observed maximum composite likelihood where the expected SFS is replaced by the relative observed SFS during the calculation of fastsimcoal2, and CLe is the estimated composite maximum likelihood. The null distribution of CLR is obtained from the same simulated data sets used to infer the confidence interval above, where each dataset can generate a CLR statistic. The p-value of the chosen final demographic parameters can be then be calculated from how many CLR statistics are larger than the chosen model’s CLR statistic. If the p-value is not significant, it indicates that the observed SFS is well explained by the chosen model.

## Accession Number

All the sequences have been deposited in GSA (The Genome Sequence Archive database, http://gsa.big.ac.cn/) with Accession ID (PRJCA000251).

## Acknowledgments

This work was supported by the Strategic Priority Research Program of the Chinese Academy of Sciences, Grant No XDB13020600. We thank the collaborators who kindly shared brown rat samples: J.-M. Duplantier, L. Granjon, K. Bâ, C. Brouat, M. Diallo, M. Kane, A. Sow (IRD, CBGP), B. Sicard (IRD, IMBE), S. Traoré (Institut d’Economie Rurale), C. Goarant (Institut Pasteur de Nouvelle Calédonie), S. Piry, Y. Chaval, N. Charbonnel (INRA, CBGP), C. Gotteland (CNRS/Université Lyon 1), Alexey P. Kryukov, Mr. Erwan Lagadec and Dr Pablo Tortosa (CRVOI-PIMIT). Authorization to use rat samples from the Seychelles collected during a CRVOI-IRD mission (SBS authorization dated 24.05.11) was provided by the Ministry of the Environment, Energy and Climate Change of Seychelles (special thanks to M. Alain de Comarmond, Principal Secretary, and M. Ronley Fanchette, Director of Conservation). Samples hosted at CBGP are stored at the collection platform (http://www6.montpellier.inra.fr/cbgp_eng/Platforms/Collections-platform), and included in the small mammal database (http://vminfotron-dev.mpl.ird.fr/bdrss/index.php).

